# Transcript Assembly and Annotations: Bias and Adjustment

**DOI:** 10.1101/2023.04.20.537700

**Authors:** Qimin Zhang, Mingfu Shao

**Affiliations:** Department of Computer Science and Engineering, School of Electrical Engineering and Computer Science, The Pennsylvania State University; Huck Institutes of the Life Sciences, The Pennsylvania State University

**Keywords:** transcript annotation, transcript assembly, intron retention

## Abstract

**Motivation:** Transcript annotations play a critical role in gene expression analysis as they serve as a reference for quantifying isoform-level expression. The two main sources of annotations are RefSeq and Ensembl/GENCODE, but discrepancies between their methodologies and information resources can lead to significant differences. It has been demonstrated that the choice of annotation can have a significant impact on gene expression analysis. Furthermore, transcript assembly is closely linked to annotations, as assembling large-scale available RNA-seq data is an effective data-driven way to construct annotations, and annotations are often served as benchmarks to evaluate the accuracy of assembly methods. However, the influence of different annotations on transcript assembly is not yet fully understood.

**Results:** We investigate the impact of annotations on transcript assembly. We observe that conflicting conclusions can arise when evaluating assemblers with different annotations. To understand this striking phenomenon, we compare the structural similarity of annotations at various levels and find that the primary structural difference across annotations occurs at the intron-chain level. Next, we examine the biotypes of annotated and assembled transcripts and uncover a significant bias towards annotating and assembling transcripts with intron retentions, which explains above the contradictory conclusions. We develop a standalone tool, available at https://github.com/Shao-Group/irtool, that can be combined with an assembler to generate an assembly without intron retentions. We evaluate the performance of such a pipeline and offer guidance to select appropriate assembling tools for different application scenarios.

## 1 Introduction

The isoform-level expression analysis has become a common toolbox in biological and biomedical studies. This analysis generally involves quantifying the expression levels of annotated transcripts given the RNA-seq data, followed by statistical methods to identify differentially expressed transcripts. Popular tools used in this pipeline include RSEM [1], kallisto [2], Salmon [3], edgeR [4], and DESeq2 [5], among others. The transcriptome, which is the collection of annotated transcripts, plays a crucial role in the analysis, as it serves as a reference for isoform quantification and splicing quantification [6, 7, 8]. The construction of high-quality, reliable, and complete transcriptomes for model species has been a long-standing community effort. Currently, RefSeq [9], led by NCBI, and Ensembl/GENCODE [10], led by EMBL-EBI, are the two main sources of annotations, with updates and curations constantly being made.

It is widely acknowledged that the RefSeq and Ensembl annotations differ significantly due to differences in methodology and information resources. Generally, RefSeq annotation prioritizes experimental evidence, while Ensembl annotation incorporates more computational predictions and includes more novel splicing variants [11]. The choice of an annotation depends on the specific need; for example, RefSeq is commonly used for variant studies [12], while Ensembl annotations are preferred for extensive research initiatives such as ENCODE [13], gnomAD [14], and GTEx [15]. Several studies have investigated the impact of different annotations on gene expression analysis. It has been reported that the choice of annotation has a significant effect on RNA-seq read mapping, gene/isoform quantification, and differential analysis [16, 17, 18]. In addition, integrating diverse annotations can markedly improve transcriptomic and genetic studies [19].

Computational methods are increasingly used to identify novel isoforms to complement annotations of model species and to construct transcriptomes for non-model species, thanks to the availability of large-scale reposited RNA-seq data. The process of reconstructing full-length transcripts from RNA-seq reads, known as transcript assembly, has been extensively studied, with significant progress made in advancing the theory [20, 21, 22] and in developing practical assemblers, including Cufflinks [23], CLASS2 [24], TransComb [25], FLAIR [26], StringTie [27], Scallop [28], StringTie2 [29], and Scallop2 [30], to just name a few. There is a close relationship between transcript assembly and annotations. On one hand, transcript assembly offers a data-driven venue to construct annotations [31]; on the other hand, annotations serve as a ground-truth to evaluate assemblers on real RNA-seq data where the true expressed transcripts are unknown. We study the impact of annotations on transcript assembly in this work. We evaluate the accuracy of two recent and popular assemblers Scallop2 and StringTie2 with different annotations (Section 2.1). Surprisingly, we found that Scallop2 performs better than StringTie2 when evaluated with Ensembl annotations but worse with RefSeq annotations. To uncover the underlying reasons, we first systematically compare the structural similarity between different annotations to investigate the primary sources of divergence (Section 2.2). We found that while annotations already differ significantly at the intron-exon boundary and junction levels, the differences are most pronounced at the intron-chain level. We then investigate if the differences are related to transcript biotypes (Section 2.3). We observed that transcripts with intron retentions contribute the most significant disparity to RefSeq and Ensembl annotations. Meanwhile, Scallop2 and StringTie2 also behave differently in assembling such transcripts. We therefore conclude that the joint bias in assembling and annotating transcripts with intron retentions leads to the opposite evaluation results. Finally, we propose criteria and develop a tool to adjust the biases in intron retention for an assembly and provide guidance for a suitable pipeline based on testing the assemblers with and without using the adjustment (Section 2.4).

## 2 Results

### 2.1 Evaluating Assemblers with Different Annotations

We show that divergent conclusions can be drawn when transcript assemblers are evaluated using different annotations. We investigate this phenomenon by comparing two widely-used reference-based assemblers, StringTie2 and Scallop2, with five transcriptome annotations derived from two human genome build, GRCh38 and T2T-CHM13 [32], on 17 paired-end RNA-seq samples from two datasets, 10 samples from EN10 and 7 samples from HS7, aligned with two popular aligners, STAR [33] and HISAT2 [34]. The assembly accuracies are evaluated using GffCompare [35]. Details about the comparison of the methods, accuracy measures, and evaluation pipeline are provided in Section 4.1. We do not intend to conduct a comprehensive benchmarking analysis for assemblers but rather to reveal the divergence of annotations and their impacts on evaluating assemblers.

Table 1 compares the accuracy of StringTie2 and Scallop2. As shown in the table, Scallop2 outperforms StringTie2 when evaluated with Ensembl and CHM13 annotations, evidenced by Scallop2 outperforming on all samples (68 and 34, respectively; using adjusted precisions to break ties). However, a different conclusion is reached when evaluated with RefSeq annotations, as StringTie2 outperforms on more samples (38 out of 68) than Scallop2. The high level of agreement between Ensembl and CHM13 annotations (from T2T-CHM13 genome build) is expected since they are very similar, as illustrated in Figures 3 and 4. We therefore focus on exploring the differences between Ensembl and RefSeq annotations.

**Table 1:**
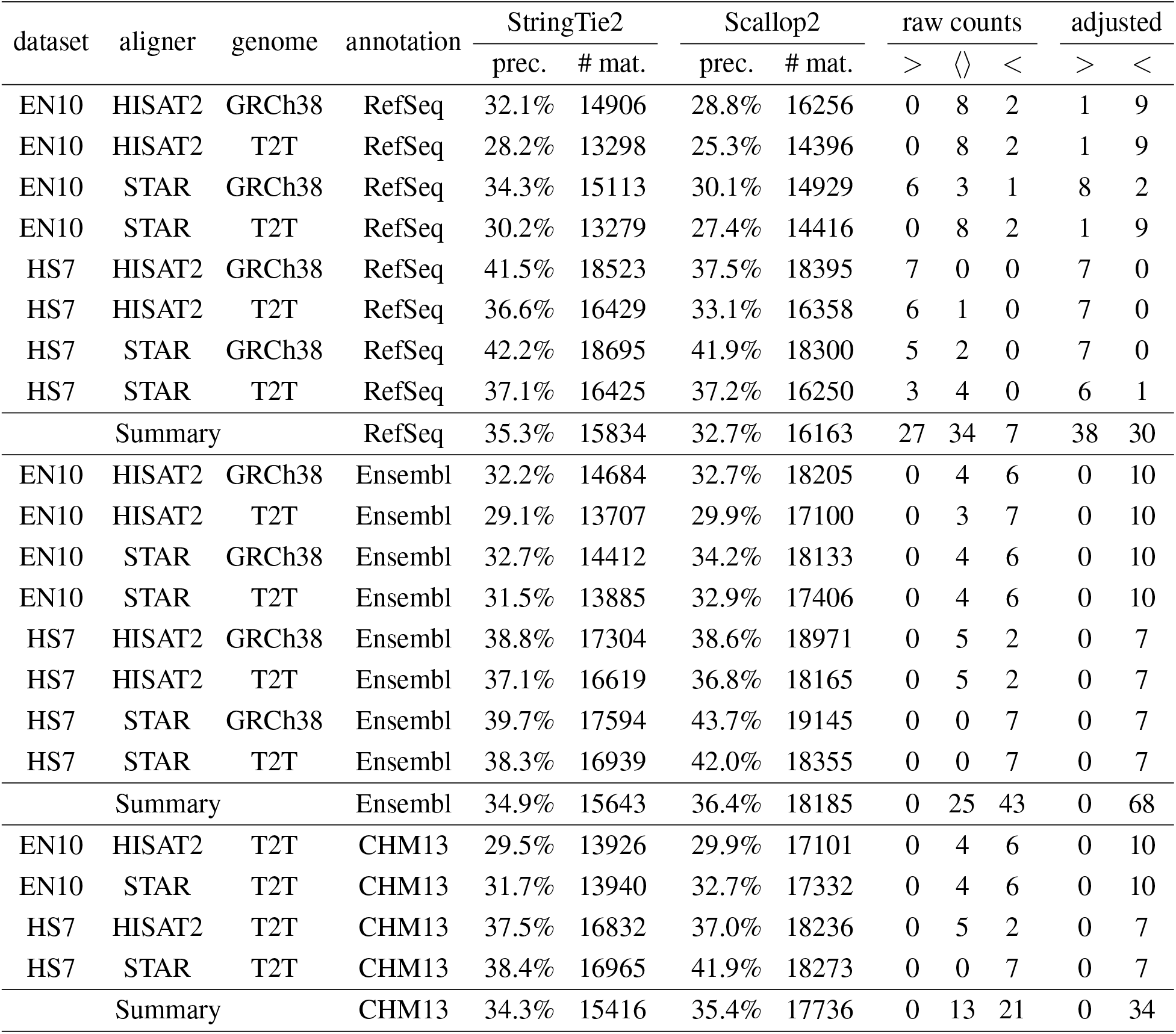
Comparison of the assembly accuracy, measured with precision (%) and the number of matching transcripts, of StringTie2 and Scallop2 using different annotations as the reference. In each combination (of dataset, aligner, genome build, annotation) the two metrics are averaged over all samples in the dataset. Symbol ⟨⟩ indicates that one method gets higher on one metric but lower on the other; symbol *>* indicates that StringTie2 outperforms on both metrics, while *<* indicates Scallop2 outperforms on both metrics. The three columns of *raw counts* give the number of samples in each category by comparing raw precision and recall. Samples in the ⟨⟩ category are further compared using the adjusted precision, and the number of samples are merged into either *>* or *<* category accordingly, shown in the two columns of under *adjusted*.

| dataset | aligner | genome | annotation | StringTie2 |  | Scallop2 |  | raw counts |  |  | adjusted |  |
| --- | --- | --- | --- | --- | --- | --- | --- | --- | --- | --- | --- | --- |
| | | | | prec. | # mat. | prec. | # mat. | $>$ | $\langle \rangle$ | $<$ | $>$ | $<$ |
| EN10 | HISAT2 | GRCh38 | RefSeq | 32.1% | 14906 | 28.8% | 16256 | 0 | 8 | 2 | 1 | 9 |
| EN10 | HISAT2 | T2T | RefSeq | 28.2% | 13298 | 25.3% | 14396 | 0 | 8 | 2 | 1 | 9 |
| EN10 | STAR | GRCh38 | RefSeq | 34.3% | 15113 | 30.1% | 14929 | 6 | 3 | 1 | 8 | 2 |
| EN10 | STAR | T2T | RefSeq | 30.2% | 13279 | 27.4% | 14416 | 0 | 8 | 2 | 1 | 9 |
| HS7 | HISAT2 | GRCh38 | RefSeq | 41.5% | 18523 | 37.5% | 18395 | 7 | 0 | 0 | 7 | 0 |
| HS7 | HISAT2 | T2T | RefSeq | 36.6% | 16429 | 33.1% | 16358 | 6 | 1 | 0 | 7 | 0 |
| HS7 | STAR | GRCh38 | RefSeq | 42.2% | 18695 | 41.9% | 18300 | 5 | 2 | 0 | 7 | 0 |
| HS7 | STAR | T2T | RefSeq | 37.1% | 16425 | 37.2% | 16250 | 3 | 4 | 0 | 6 | 1 |
| Summary |  |  | RefSeq | 35.3% | 15834 | 32.7% | 16163 | 27 | 34 | 7 | 38 | 30 |
| EN10 | HISAT2 | GRCh38 | Ensembl | 32.2% | 14684 | 32.7% | 18205 | 0 | 4 | 6 | 0 | 10 |
| EN10 | HISAT2 | T2T | Ensembl | 29.1% | 13707 | 29.9% | 17100 | 0 | 3 | 7 | 0 | 10 |
| EN10 | STAR | GRCh38 | Ensembl | 32.7% | 14412 | 34.2% | 18133 | 0 | 4 | 6 | 0 | 10 |
| EN10 | STAR | T2T | Ensembl | 31.5% | 13885 | 32.9% | 17406 | 0 | 4 | 6 | 0 | 10 |
| HS7 | HISAT2 | GRCh38 | Ensembl | 38.8% | 17304 | 38.6% | 18971 | 0 | 5 | 2 | 0 | 7 |
| HS7 | HISAT2 | T2T | Ensembl | 37.1% | 16619 | 36.8% | 18165 | 0 | 5 | 2 | 0 | 7 |
| HS7 | STAR | GRCh38 | Ensembl | 39.7% | 17594 | 43.7% | 19145 | 0 | 0 | 7 | 0 | 7 |
| HS7 | STAR | T2T | Ensembl | 38.3% | 16939 | 42.0% | 18355 | 0 | 0 | 7 | 0 | 7 |
| Summary |  |  | Ensembl | 34.9% | 15643 | 36.4% | 18185 | 0 | 25 | 43 | 0 | 68 |
| EN10 | HISAT2 | T2T | CHM13 | 29.5% | 13926 | 29.9% | 17101 | 0 | 4 | 6 | 0 | 10 |
| EN10 | STAR | T2T | CHM13 | 31.7% | 13940 | 32.7% | 17332 | 0 | 4 | 6 | 0 | 10 |
| HS7 | HISAT2 | T2T | CHM13 | 37.5% | 16832 | 37.0% | 18236 | 0 | 5 | 2 | 0 | 7 |
| HS7 | STAR | T2T | CHM13 | 38.4% | 16965 | 41.9% | 18273 | 0 | 0 | 7 | 0 | 7 |
| Summary |  |  | CHM13 | 34.3% | 15416 | 35.4% | 17736 | 0 | 13 | 21 | 0 | 34 |

We further demonstrate the dramatic discrepancy of RefSeq and Ensembl annotations in evaluating transcript assemblers, by comparing their accuracies on individual samples. The results are shown in Figure 1 (using GRCh38 annotations) and Figure 2 (using T2T-CHM13 annotations). Across all combinations, Scallop2’s accuracy is significantly higher under Ensembl annotation than under RefSeq. Conversely, the pattern is almost reversed for StringTie2, with its accuracy evaluated under Ensembl being either lower than that under RefSeq in the case of GRCh38 annotations (Figure 1), or only slightly higher in the case of T2T-CHM13 annotations (Figure 2).

**Figure 1:**
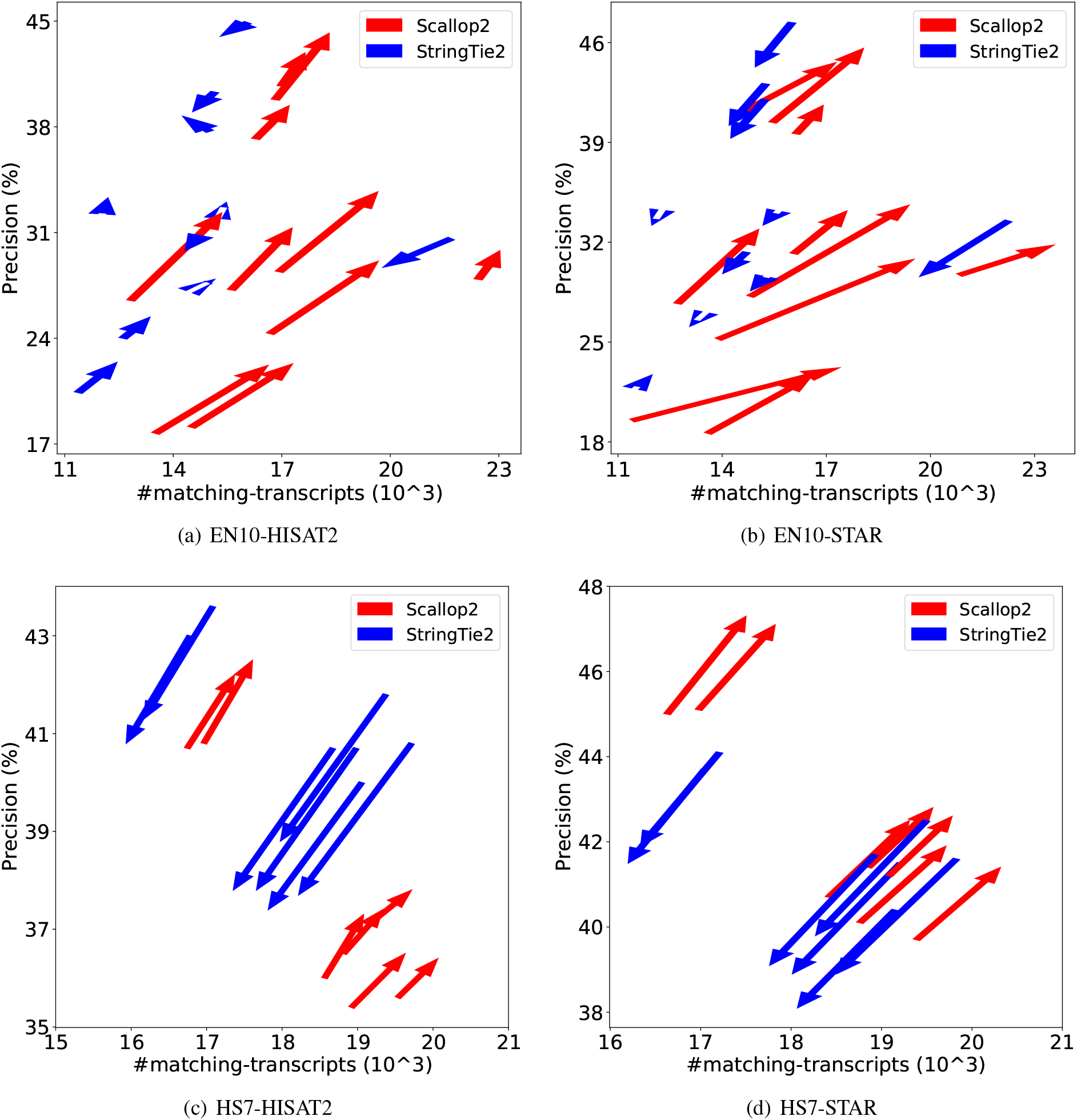
Illustrating the difference of assembly accuracy evaluated with RefSeq and Ensembl annotations, both from GRCh38 genome build. Each arrow represents a sample, pointing from the accuracy evaluated with RefSeq annotation to that with Ensembl annotation. The subfigures correspond to the four combinations of dataset (EN10 or HS7) and aligner (HISAT2 or STAR).

**Figure 2:**
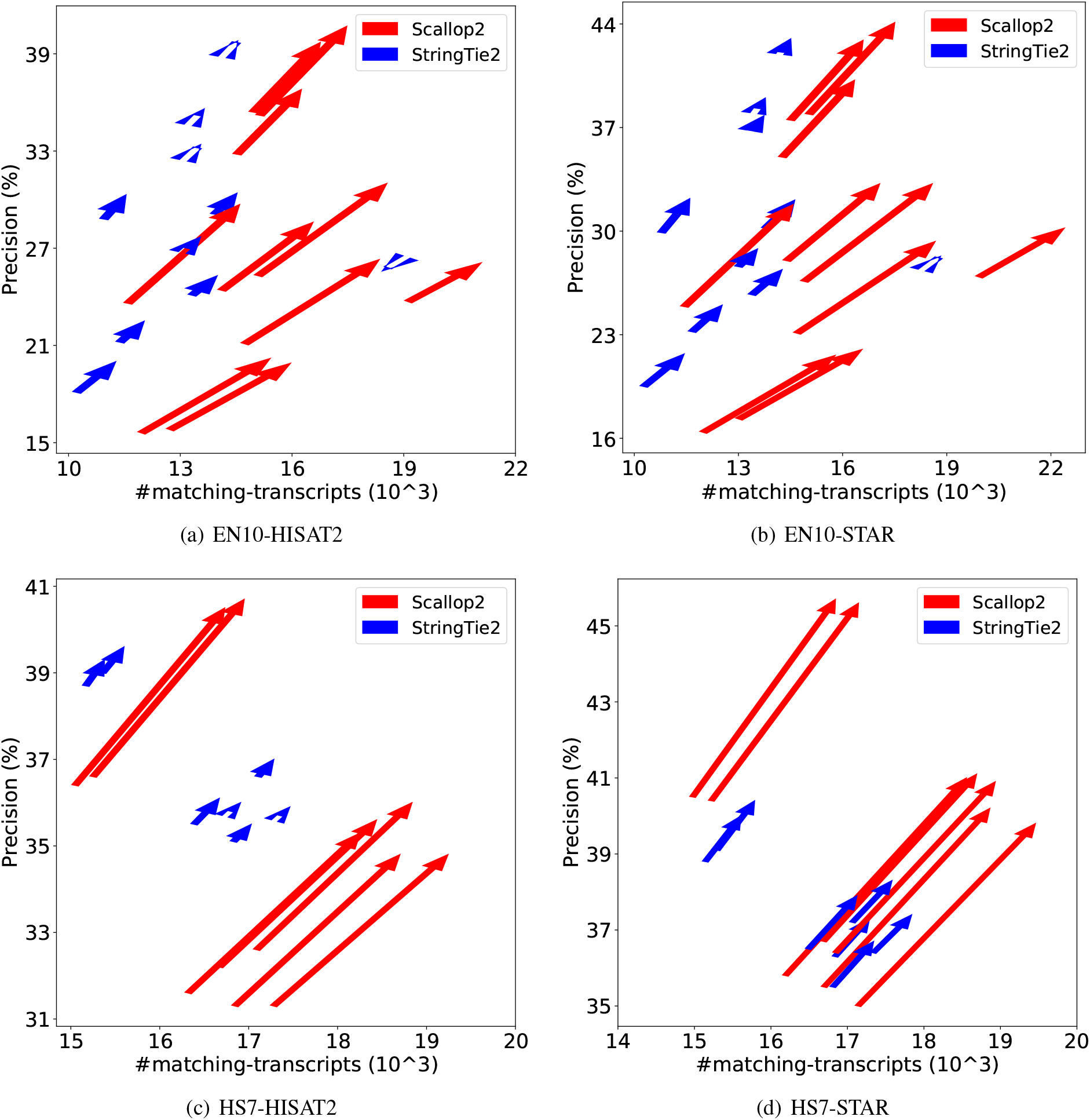
Illustrating the difference of assembly accuracy evaluated with RefSeq and Ensembl annotations, both from T2T-CHM13 genome build. Each arrow represents a sample, pointing from the accuracy evaluated with RefSeq annotation to that with Ensembl annotation. The subfigures correspond to the four combinations of dataset (EN10 or HS7) and aligner (HISAT2 or STAR).

### 2.2 Comparison of Structural Similarities

The results presented in Section 2.1 clearly highlight the substantial differences between RefSeq and Ensembl annotations. This prompts us to investigate the primary sources of these divergences. In particular, since a transcript can be represented as a chain of alternating exons and introns, we seek to determine the level of transcript structure that contributes the most to these differences, whether it be at the individual exon-intron boundary, the junction (pair of intron boundaries), or the chain of junctions. To address this question, we propose several metrics and use them to evaluate the similarity of annotations at these three levels. Details about the metrics definitions are provided in Section 4.2.

We plotted the Jaccard similarities across different annotations in Figure 3. It clearly shows that the Ensembl and CHM13 annotations of the T2T-CHM13 genome build are highly similar, with Jaccard of 0.91 at the boundary and junction levels, and 0.80 at the intron-chain level. However, the Ensembl and RefSeq annotations in both genome builds exhibit significant divergence, with Jaccard values lower than 0.69 and 0.57 at the boundary and junction levels, respectively. This difference is most pronounced at the intron-chain level, where the Jaccard similarity drops to 0.19, indicating that the intron-chain is the primary contributor to the structural disparity between Ensembl and RefSeq annotations.

**Figure 3:**
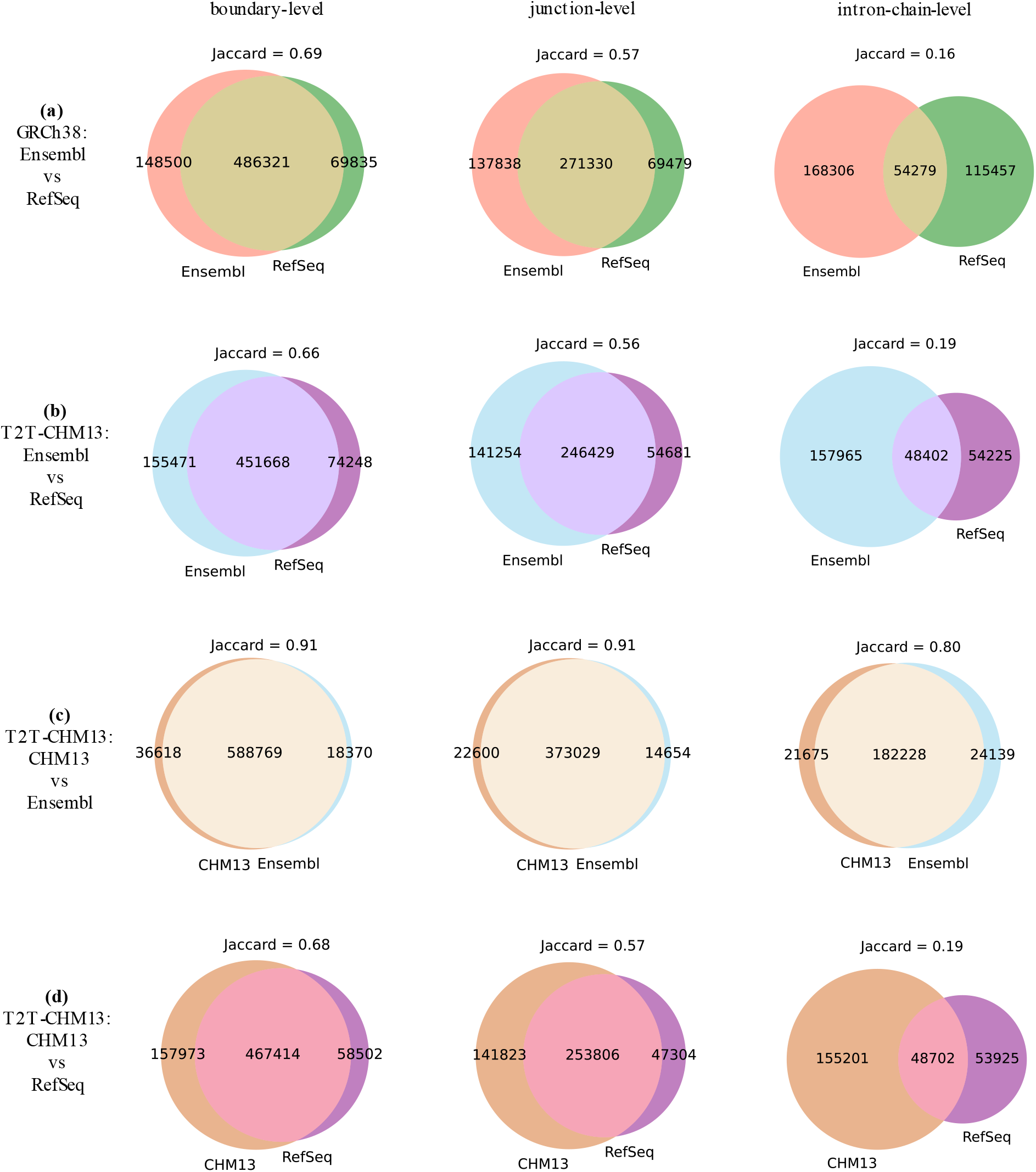
The Jaccard similarity of 4 pairs of annotations (4 subfigures) at the level of boundary, junction, and intron-chain.

Next, we measure the similarity of individual genes in different annotations, aiming to determine whether the difference of annotations can be attributed to a small portion of genes. The method to construct the correspondence between genes in two annotations and definitions of Jaccard similarities of every constructed gene pair at boundary, junction, and intron-chain levels, are provided in Section 4.3. The gene pairs constructed with this approach aligns very well with gene nomenclature. Specifically, there are 24957 gene pairs between RefSeq and Ensembl annotations according to the HUGO Gene Nomenclature. Among these, 23478 pairs (94.1%) can be found in the gene pairs constructed with our approach.

The distribution of the Jaccard similarities are shown in Figure 4. As before, we observe that Ensembl and CHM13 annotations in the T2T-CHM13 genome build are almost identical to each other. However, Ensembl and RefSeq annotations in either genome build show significant divergence, especially at the intron-chain level. Furthermore, the majority of the gene pairs between Ensembl and RefSeq annotations are divergent, as evidenced by the quartiles in Figure 4.

**Figure 4:**
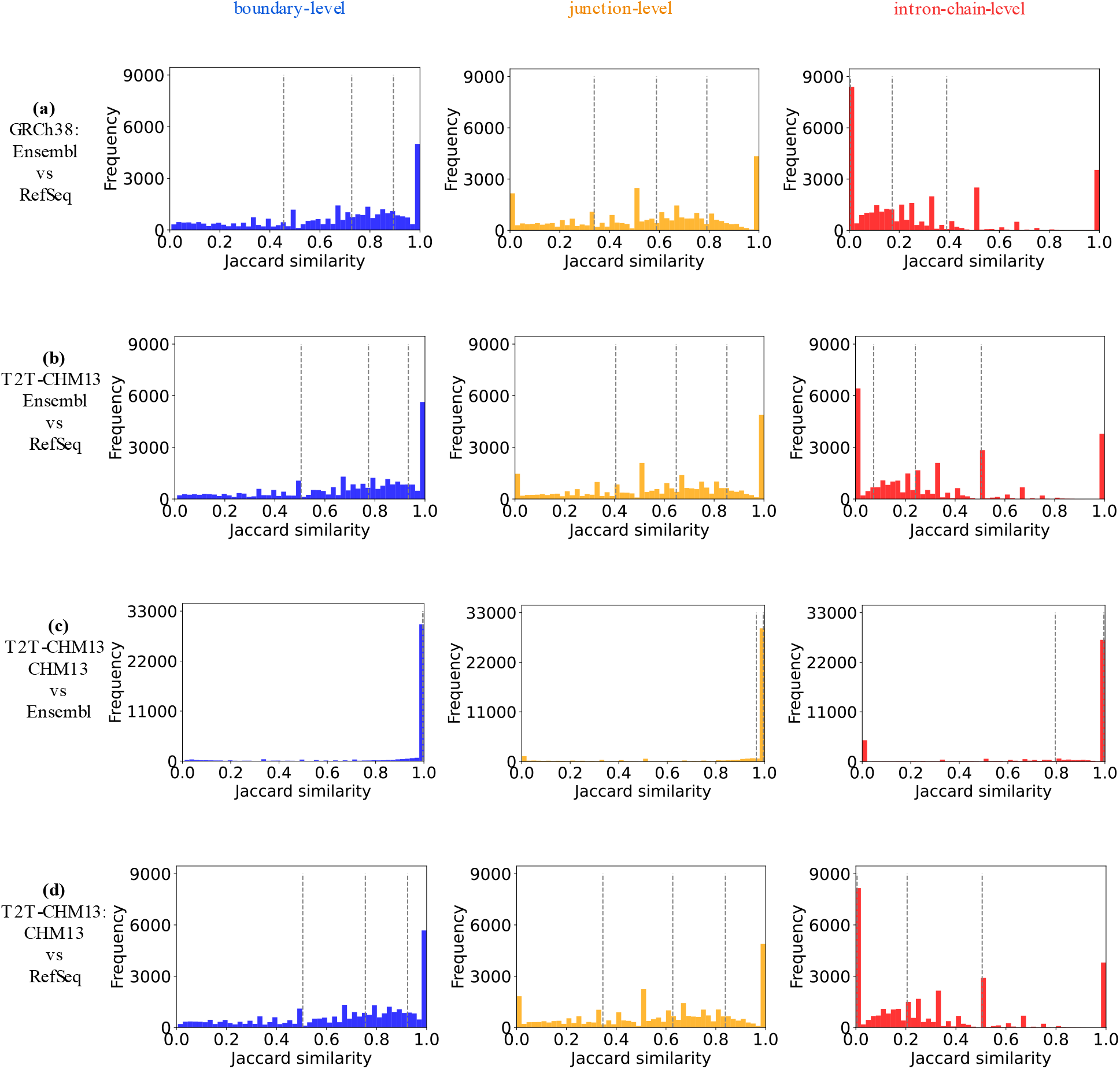
The distribution of Jaccard similarities of all paired genes in each pair of compared annotations at the level of boundary, junction, and intron-chain. The three dashed lines in each subfigure mark the Jaccard similarity at the 25th, 50th, and 75th percentile of the total frequency.

### 2.3 Comparison of Transcript Biotypes

The Ensembl annotation provides each annotated transcript with a “biotype” that indicates its biological category and function. By leveraging this information, we aim to investigate whether different annotations or assemblers show bias towards specific biotypes.

In Table 2, we compare the distribution of multi-exon transcripts belonging to different biotypes between the Ensembl and RefSeq annotations. As biotype information is not available in the RefSeq annotation, we report the number and percentage of transcripts annotated by Ensembl in each biotype that are also annotated in RefSeq. (Two transcripts are considered the same if they share the same intron-chain.) Our analysis reveals huge divergences in several biotypes, such as “retained intron”, “processed transcript”, and “processed pseudogene”, where only a tiny portion of them are annotated in RefSeq. Of particular interest is the “retained intron” biotype, which is the third largest biotype in the Ensembl annotation, but only 1.3% of them appear in the RefSeq annotation.

**Table 2:**
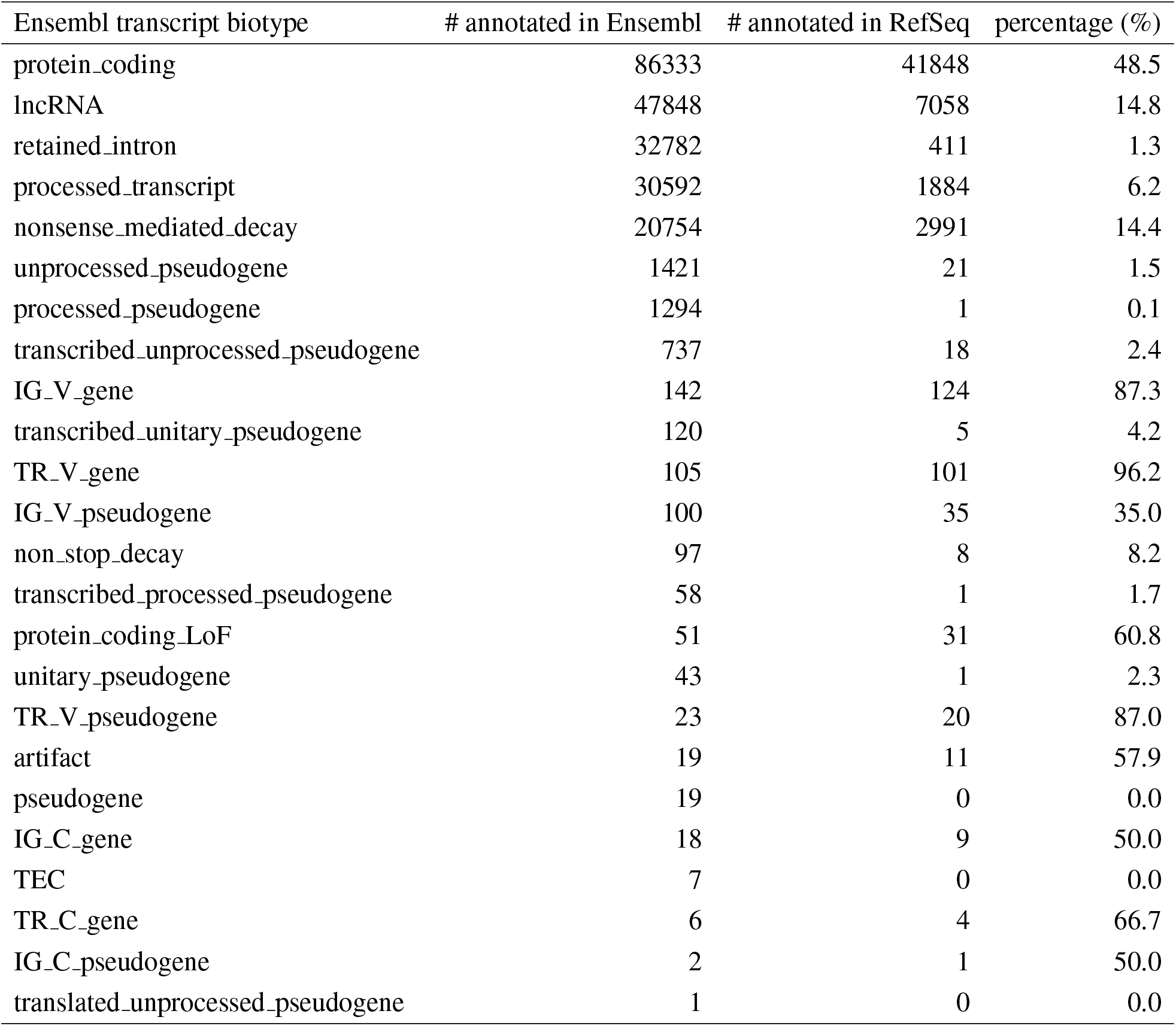
Illustration of the number of multi-exon transcripts in different biotypes between Ensembl and RefSeq annotations (GRCh38 build). The first column lists biotypes defined by the Ensembl annotation; the second column lists the number of multi-exon transcripts in each biotype; the third and the fourth columns give the number and the percentage of multi-exon transcripts in each biotype that are also annotated in the RefSeq annotation.

We investigate whether the dissimilarities between Scallop2 and StringTie2 when evaluated with Ensembl and RefSeq annotations (as discussed in Section 2.1) can be attributed to differences in biotypes. We classify the matching transcripts based on their biotypes, using the Ensembl annotation as the reference. The comparison of the five largest biotypes is presented in Figure 5. Notably, Scallop2 identifies significantly more matching transcripts in the “retained intron” biotype compared to StringTie2. We therefore conjecture that it is the bias towards retained-intron transcripts, i.e., Scallop2 assembles more such transcripts than StringTie2 while Ensembl annotates more such transcripts than RefSeq, that is the primary reason behind the opposite conclusion when StringTie2 and Scallop2 are evaluated against the two annotations.

**Figure 5:**
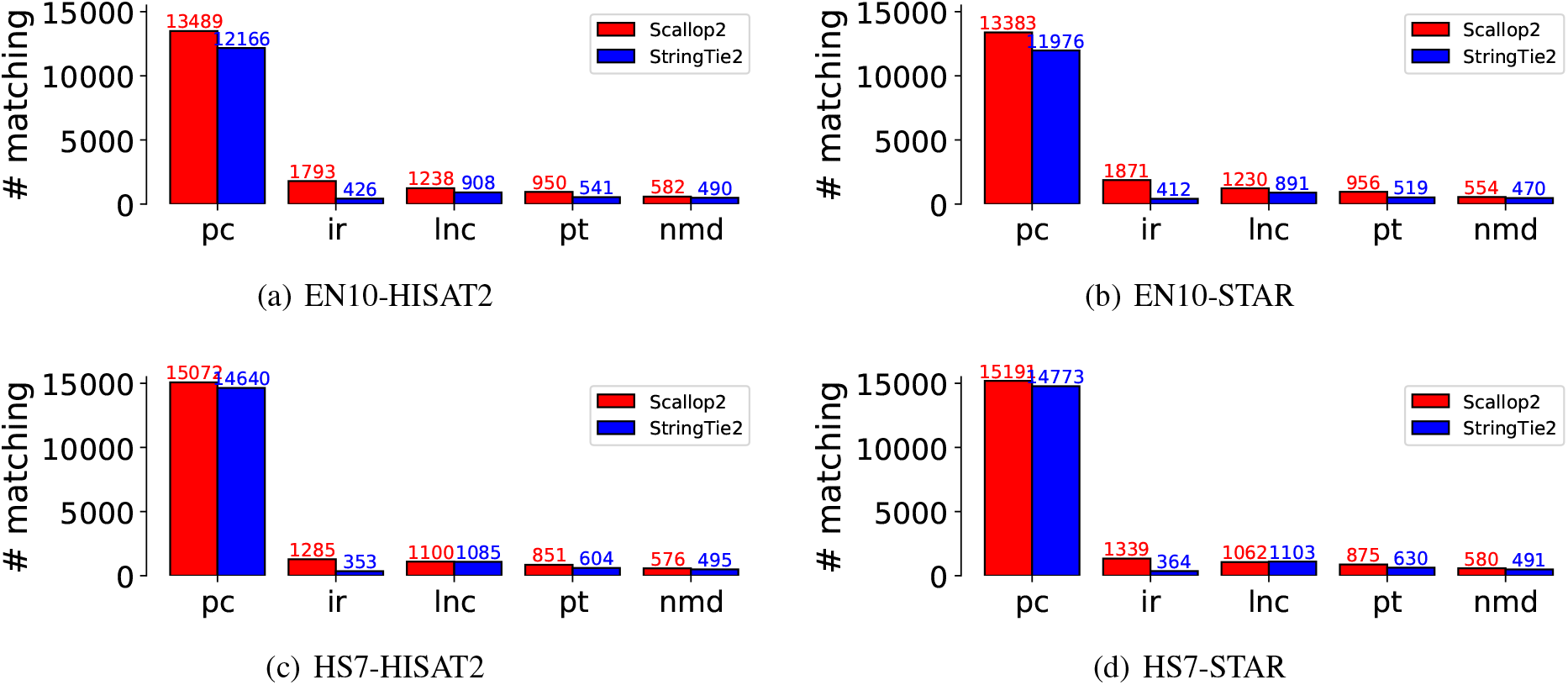
Comparison of the matching transcripts assembled by StringTie2 and Scallop2 in the five largest biotypes using the Ensembl annotation as the reference. The average number over all samples in each dataset is reported in the barplot. The 5 biotypes are: protein coding (pc), retained intron (ir), lncRNA (lnc), processed transcript (pt), and nonsense mediated decay (nmd). These five biotypes account for 98.6% of the total transcripts in Ensembl.

To further support the above hypothesis, we analyzed the number of assembled transcripts that were annotated in Ensembl but not in RefSeq. These transcripts are considered true positives by Ensembl and false positives by RefSeq. The results for the five largest biotypes are presented in Figure 6, which shows that the largest difference is due to the “retained intron” biotype. Therefore, we conclude that the discrepancy in evaluation between Scallop2 and StringTie2 is caused by the differences in transcripts with intron retentions between Ensembl and RefSeq annotations.

**Figure 6:**
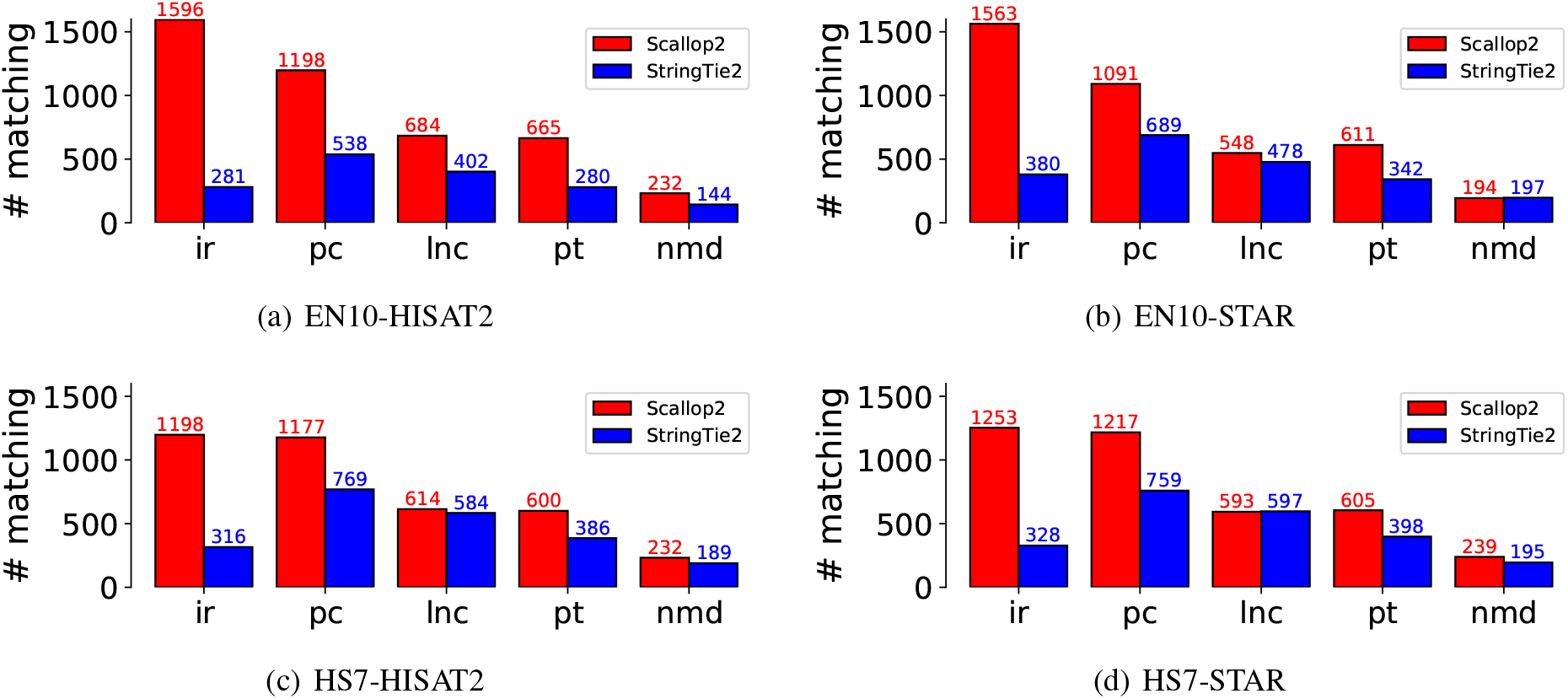
Comparison of the transcripts assembled by Scallop2 and StringTie2 that are annotated by Ensembl but not by RefSeq in each of the five largest biotypes. The average number over all samples in each dataset is reported in the barplot. The 5 biotypes are ir: retained intron, pc: protein coding, lnc: lncRNA (long non-coding RNA), pt: processed transcript, nmd: nonsense mediated decay.

### 2.4 Quantifying the Impact of Intron Retentions

We hence investigate intron retentions. We first establish a formal definition for (partial) intron retentions in the context of assembly (see Section 4.4). We then develop a tool, namely irtool, based on this definition that can identify transcripts with intron retentions. This tool can be applied to an assembly generated by any assembler, allowing for the extraction or filtering out of transcripts with retained introns. Using this tool, we quantify the impact of intron retention on Scallop2 and StringTie2.

We apply irtool to filter out transcripts with (partial) intron retentions in the StringTie2 and Scallop2 assemblies, and then evaluate the filtered assemblies using RefSeq and Ensembl annotations. Based on the fact that Ensembl annotates many more transcripts with intron retention than RefSeq (Table 2), and that Scallop2 assembles more such transcripts than StringTie2 (Figures 5 and 6), we expect that the accuracy of Scallop2 would decrease under Ensembl annotation and increase under RefSeq annotation after filtering. The experimental results confirm this conjecture, as shown in Table 3. When evaluated with RefSeq, Scallop2’s precision improve by 27.0% with a small decrease in recall (4.4%), while StringTie2 gains a 7.3% increase in precision but loses 4.0% in recall. When evaluated with Ensembl, Scallop2’s precision increases by 14.0%, but there is a significant (14.3%) drop in recall. In contrast, StringTie2 improves by 4.8% in precision but decreases by 6.0% in recall. The changes in accuracy resulting from filtering in individual samples are shown in Figure 7–8 (GRCh38 annotations) and Figures 9–10 (T2T-CHM13 annotations). In all cases (combinations of dataset, aligner, and genome build used), Scallop2 shows a sharper slope than StringTie2 on RefSeq annotations, indicating a much higher gain in precision with a similar decrease in recall.

**Figure 7:**
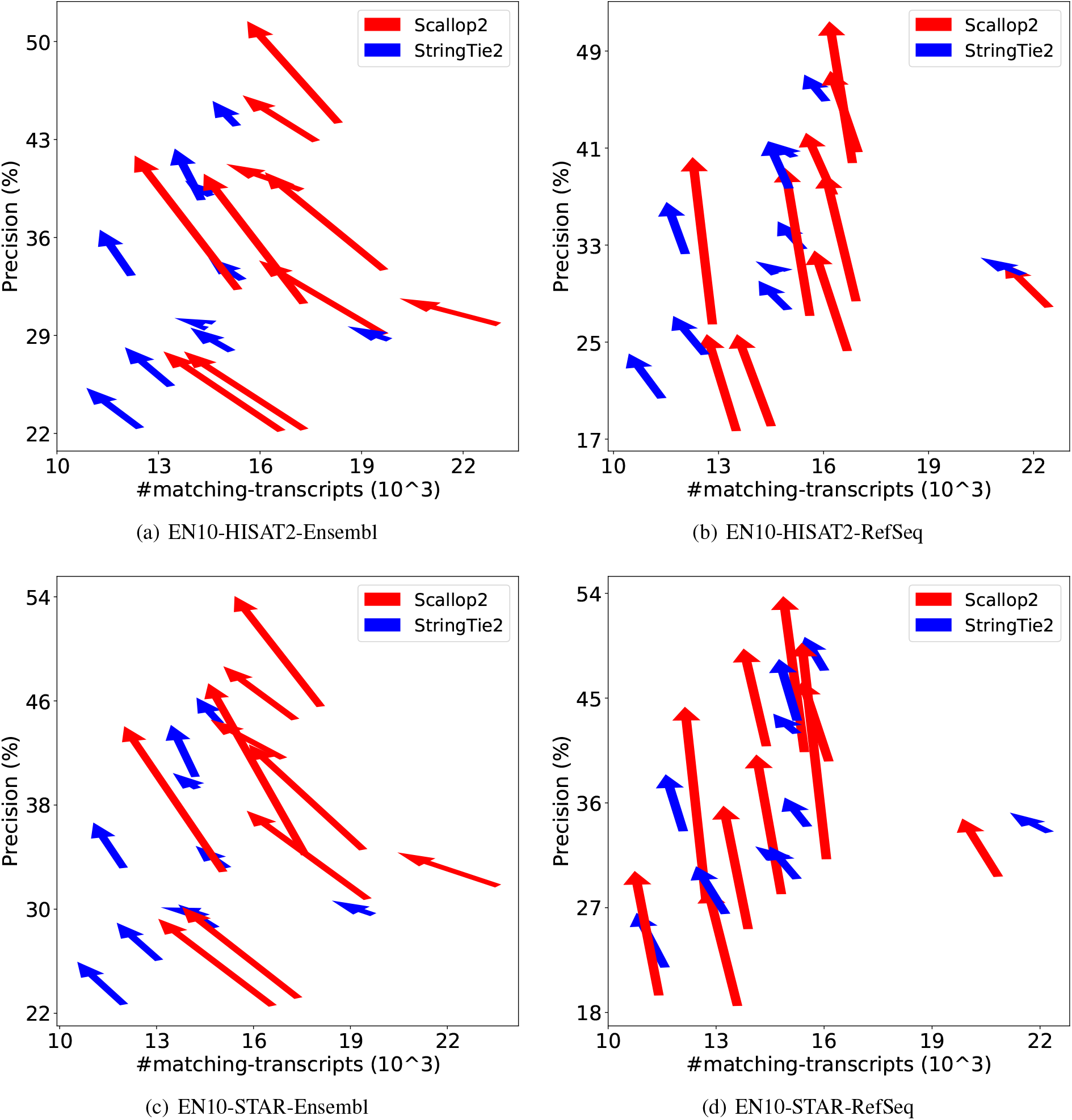
Comparison of assembly accuracy of StringTie2 and Scallop2 before and after filtering out transcripts with intron retention. Each arrow represents a sample, pointing from the accuracy before filtering to that after filtering. The subfigures correspond to the 4 combinations of aligner (HISAT2 or STAR) and annotations (RefSeq or Ensembl) tested on EN10; both annotations are from GRCh38 genome build.

**Figure 8:**
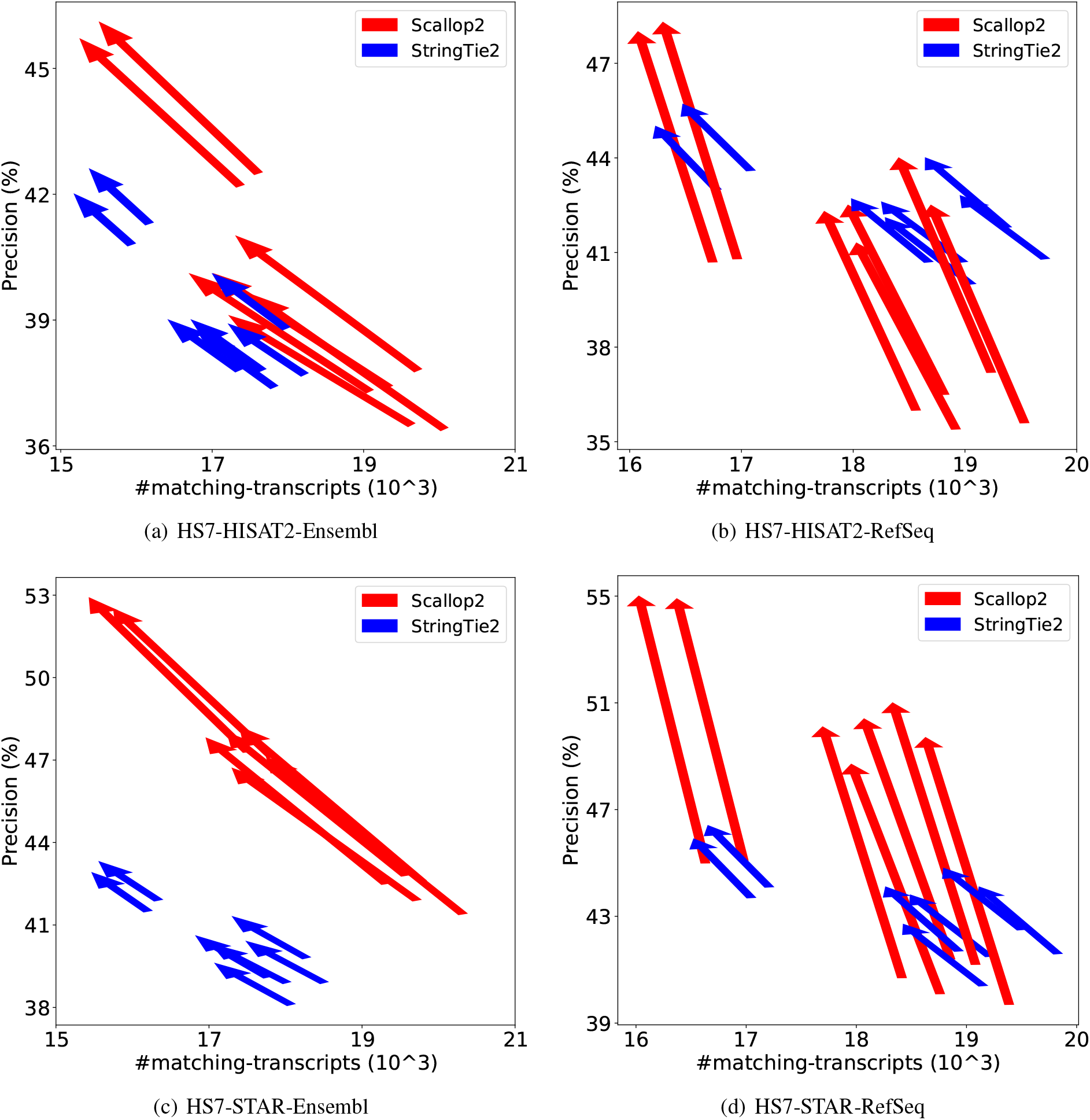
Comparison of assembly accuracy of StringTie2 and Scallop2 before and after filtering out transcripts with intron retention. Each arrow represents a sample, pointing from the accuracy before filtering to that after filtering. The subfigures correspond to the 4 combinations of aligner (HISAT2 or STAR) and annotations (RefSeq or Ensembl) tested on HS7; both annotations are from GRCh38 genome build.

**Figure 9:**
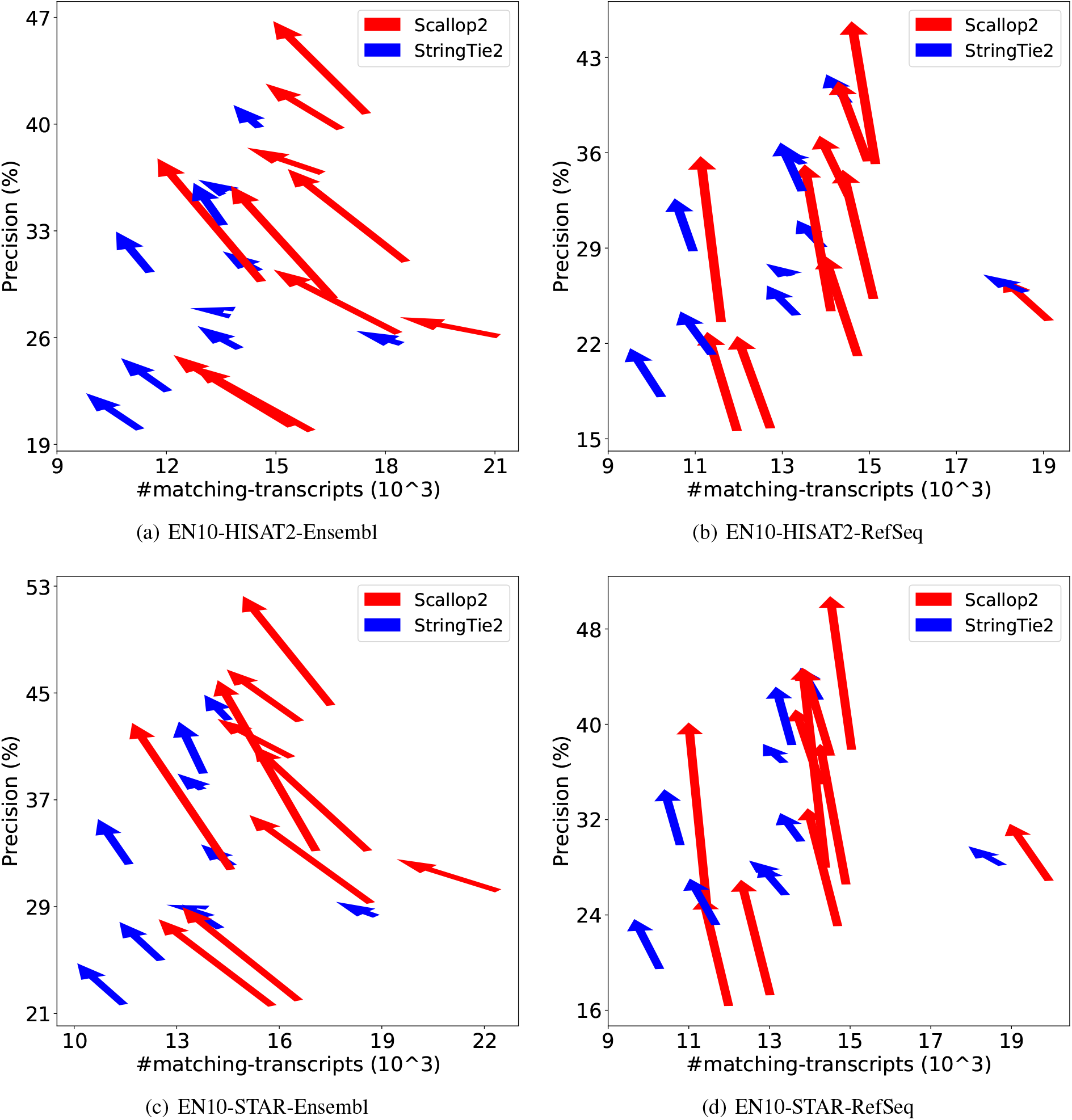
Comparison of assembly accuracy of StringTie2 and Scallop2 before and after filtering out transcripts with intron retention. Each arrow represents a sample, pointing from the accuracy before filtering to that after filtering. The subfigures correspond to the 4 combinations of aligner (HISAT2 or STAR) and annotations (RefSeq or Ensembl) tested on EN10; both annotations are from T2T-CHM13 genome build.

**Figure 10:**
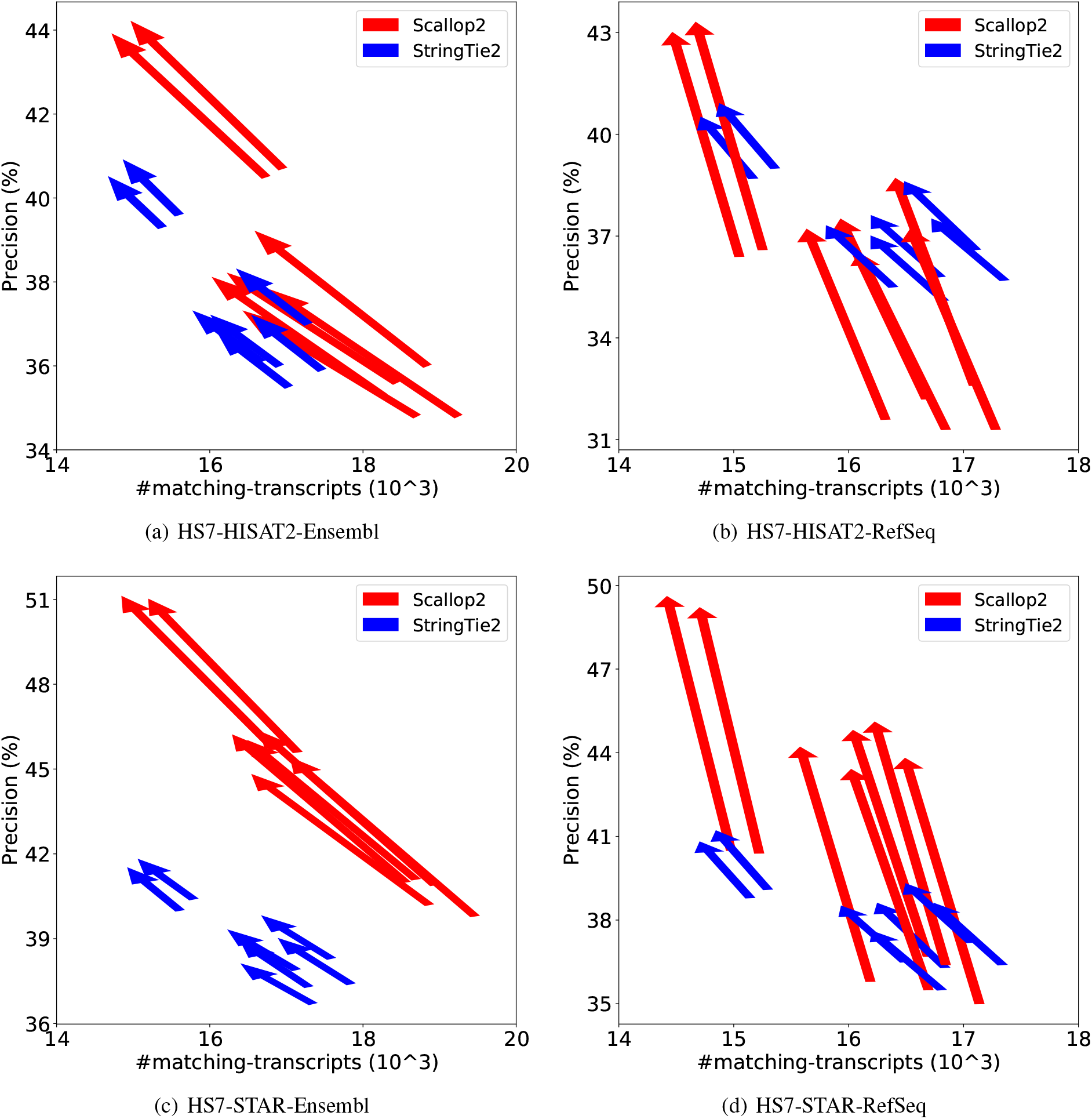
Comparison of assembly accuracy of StringTie2 and Scallop2 before and after filtering out transcripts with intron retention. Each arrow represents a sample, pointing from the accuracy before filtering to that after filtering. The subfigures correspond to the 4 combinations of aligner (HISAT2 or STAR) and annotations (RefSeq or Ensembl) tested on HS7; both annotations are from T2T-CHM13 genome build.

**Table 3.** : Comparison of relative change in percentage of precision and the number of matching transcripts after filtering out transcripts with intron retentions in StringTie2 and Scallop2 assemblies, evaluated with different annotations as the reference. Numbers are averaged over all samples in each dataset.

| dataset | aligner | genome | annotation | StringTie2 |  | Scallop2 |  |
| --- | --- | --- | --- | --- | --- | --- | --- |
| | | | | $\Delta$ prec. | $\Delta$ # mat. | $\Delta$ prec. | $\Delta$ # mat. |
| EN10 | HISAT2 | GRCh38 | RefSeq | +10.7% | -5.1% | +30.1% | -5.1% |
| EN10 | HISAT2 | T2T | RefSeq | +8.2% | -4.6% | +30.4% | -4.8% |
| EN10 | STAR | GRCh38 | RefSeq | +8.9% | -4.5% | +36.9% | -4.7% |
| EN10 | STAR | T2T | RefSeq | +9.5% | -4.1% | +37.6% | -4.5% |
| HS7 | HISAT2 | GRCh38 | RefSeq | +4.9% | -3.7% | +17.8% | -4.3% |
| HS7 | HISAT2 | T2T | RefSeq | +5.1% | -3.5% | +17.8% | -4.2% |
| HS7 | STAR | GRCh38 | RefSeq | +5.5% | -3.5% | +22.7% | -3.9% |
| HS7 | STAR | T2T | RefSeq | +5.7% | -3.3% | +23.0% | -3.8% |
| Summary |  |  | RefSeq | +7.3% | -4.0% | +27.0% | -4.4% |
| EN10 | HISAT2 | GRCh38 | Ensembl | +6.0% | -6.8% | +17.1% | -16.4% |
| EN10 | HISAT2 | T2T | Ensembl | +5.3% | -7.2% | +15.4% | -16.2% |
| EN10 | STAR | GRCh38 | Ensembl | +6.4% | -6.8% | +20.5% | -16.7% |
| EN10 | STAR | T2T | Ensembl | +6.4% | -6.7% | +20.8% | -16.6% |
| HS7 | HISAT2 | GRCh38 | Ensembl | +3.2% | -5.3% | +8.0% | -12.2% |
| HS7 | HISAT2 | T2T | Ensembl | +3.4% | -5.1% | +8.1% | -12.1% |
| HS7 | STAR | GRCh38 | Ensembl | +3.7% | -5.2% | +12.3% | -12.1% |
| HS7 | STAR | T2T | Ensembl | +3.9% | -5.0% | +12.4% | -11.9% |
| Summary |  |  | Ensembl | +4.8% | -6.0% | +14.3% | -14.3% |

As observed in Section 2.1, Scallop2 performs better than StringTie2 with Ensembl annotations but worse with RefSeq annotations. We now compare the accuracy of Scallop2 after filtering (referred to as Scallop2- ft) and unfiltered StringTie2. The results, presented in Table 4, show that Scallop2-ft outperforms StringTie2 with RefSeq annotations, evidenced by its outperforming on 45 (out of 68) samples. Scallop2-ft still outperforms StringTie2 with Ensembl annotations, although the margin is not as large as the one with unfiltered Scallop2 (see Table 1). We also compare the accuracy of both assemblers after filtering, i.e., StringTie2-ft and Scallop2-ft in Table 5. The results reveal that Scallop2-ft has a draw with StringTie2-ft with RefSeq annotations, and that Scallop2-ft outperforms StringTie2-ft on all samples.

**Table 4:** Comparison of the assembly accuracy, measured with precision (%) and number of matching transcripts, of StringTie2 and Scallop2-ft using different annotations. In each combination (of dataset, aligner, genome build, annotation) the two metrics are averaged over all samples in the dataset. Symbol ⟨⟩ indicates that one method gets higher on one metric but lower on the other; symbol *>* indicates that StringTie2 outperforms on both metrics, while *<* indicates Scallop2 outperforms on both metrics. The three columns of *raw counts* give the number of samples in each category by comparing raw precision and recall. Samples in the ⟨⟩ category are further compared using the adjusted precision, and the number of samples are merged into either *>* or *<* category accordingly, shown in the two columns of under *adjusted*.

| dataset | aligner | genome | annotation | StringTie2 |  | Scallop2-ft |  | raw counts |  |  | adjusted |  |
| --- | --- | --- | --- | --- | --- | --- | --- | --- | --- | --- | --- | --- |
| | | | | prec. | # mat. | prec. | # mat. | $>$ | $\langle \rangle$ | $<$ | $>$ | $<$ |
| EN10 | HISAT2 | GRCh38 | RefSeq | 32.1% | 14906 | 37.4% | 15421 | 0 | 2 | 8 | 1 | 9 |
| EN10 | HISAT2 | T2T | RefSeq | 28.2% | 13298 | 33.0% | 13705 | 0 | 1 | 9 | 1 | 9 |
| EN10 | STAR | GRCh38 | RefSeq | 34.3% | 15113 | 41.2% | 14223 | 0 | 6 | 4 | 6 | 4 |
| EN10 | STAR | T2T | RefSeq | 30.2% | 13279 | 37.6% | 13765 | 0 | 0 | 10 | 0 | 10 |
| HS7 | HISAT2 | GRCh38 | RefSeq | 41.5% | 18523 | 44.1% | 17599 | 0 | 7 | 0 | 7 | 0 |
| HS7 | HISAT2 | T2T | RefSeq | 36.6% | 16429 | 39.1% | 15676 | 0 | 7 | 0 | 7 | 0 |
| HS7 | STAR | GRCh38 | RefSeq | 42.2% | 18695 | 51.4% | 17581 | 0 | 7 | 0 | 0 | 7 |
| HS7 | STAR | T2T | RefSeq | 37.1% | 16425 | 45.7% | 15640 | 0 | 7 | 0 | 1 | 6 |
| Summary |  |  | RefSeq | 35.3% | 15834 | 41.2% | 15451 | 0 | 37 | 31 | 23 | 45 |
| EN10 | HISAT2 | GRCh38 | Ensembl | 32.2% | 14684 | 38.3% | 15220 | 0 | 0 | 10 | 0 | 10 |
| EN10 | HISAT2 | T2T | Ensembl | 29.1% | 13707 | 34.5% | 14331 | 0 | 0 | 10 | 0 | 10 |
| EN10 | STAR | GRCh38 | Ensembl | 32.7% | 14412 | 41.2% | 15096 | 0 | 0 | 10 | 0 | 10 |
| EN10 | STAR | T2T | Ensembl | 31.5% | 13885 | 39.7% | 14518 | 0 | 0 | 10 | 0 | 10 |
| HS7 | HISAT2 | GRCh38 | Ensembl | 38.8% | 17304 | 41.7% | 16653 | 0 | 7 | 0 | 7 | 0 |
| HS7 | HISAT2 | T2T | Ensembl | 37.1% | 16619 | 39.8% | 15975 | 0 | 7 | 0 | 7 | 0 |
| HS7 | STAR | GRCh38 | Ensembl | 39.7% | 17594 | 49.0% | 16835 | 0 | 7 | 0 | 0 | 7 |
| HS7 | STAR | T2T | Ensembl | 38.3% | 16939 | 47.3% | 16162 | 0 | 7 | 0 | 0 | 7 |
| Summary |  |  | Ensembl | 34.9% | 15643 | 41.4% | 15599 | 0 | 28 | 40 | 14 | 54 |

**Table 5:** Comparison of the assembly accuracy, measured with precision (%) and number of matching transcripts, of StringTie2-ft and Scallop2-ft using different annotations. In each combination (of dataset, aligner, genome build, annotation) the two metrics are averaged over all samples in the dataset. Symbol ⟨⟩ indicates that one method gets higher on one metric but lower on the other; symbol *>* indicates that StringTie2 outperforms on both metrics, while *<* indicates Scallop2 outperforms on both metrics. The three columns of *raw counts* give the number of samples in each category by comparing raw precision and recall. Samples in the ⟨⟩ category are further compared using the adjusted precision, and the number of samples are merged into either *>* or *<* category accordingly, shown in the two columns of under *adjusted*.

| dataset | aligner | genome | annotation | StringTie2-ft |  | Scallop2-ft |  | raw counts |  |  | adjusted |  |
| --- | --- | --- | --- | --- | --- | --- | --- | --- | --- | --- | --- | --- |
| | | | | prec. | # mat. | prec. | # mat. | $>$ | $\langle \rangle$ | $<$ | $>$ | $<$ |
| EN10 | HISAT2 | GRCh38 | RefSeq | 34.6% | 14150 | 37.4% | 15421 | 0 | 3 | 7 | 1 | 9 |
| EN10 | HISAT2 | T2T | RefSeq | 30.5% | 12690 | 33.0% | 13705 | 0 | 3 | 7 | 0 | 10 |
| EN10 | STAR | GRCh38 | RefSeq | 37.3% | 14439 | 41.2% | 14223 | 2 | 4 | 4 | 6 | 4 |
| EN10 | STAR | T2T | RefSeq | 33.0% | 12738 | 37.6% | 13765 | 0 | 1 | 9 | 0 | 10 |
| HS7 | HISAT2 | GRCh38 | RefSeq | 43.6% | 17834 | 44.1% | 17599 | 4 | 3 | 0 | 7 | 0 |
| HS7 | HISAT2 | T2T | RefSeq | 38.5% | 15857 | 39.1% | 15676 | 4 | 3 | 0 | 7 | 0 |
| HS7 | STAR | GRCh38 | RefSeq | 44.5% | 18038 | 51.4% | 17581 | 0 | 7 | 0 | 0 | 7 |
| HS7 | STAR | T2T | RefSeq | 39.2% | 15883 | 45.7% | 15640 | 0 | 7 | 0 | 1 | 6 |
| Summary |  |  | RefSeq | 37.7% | 15204 | 41.2% | 15451 | 10 | 31 | 27 | 22 | 46 |
| EN10 | HISAT2 | GRCh38 | Ensembl | 34.2% | 13681 | 38.3% | 15220 | 0 | 2 | 8 | 0 | 10 |
| EN10 | HISAT2 | T2T | Ensembl | 30.6% | 12724 | 34.5% | 14331 | 0 | 2 | 8 | 0 | 10 |
| EN10 | STAR | GRCh38 | Ensembl | 34.8% | 13430 | 41.2% | 15096 | 0 | 0 | 10 | 0 | 10 |
| EN10 | STAR | T2T | Ensembl | 33.6% | 12955 | 39.7% | 14518 | 0 | 0 | 10 | 0 | 10 |
| HS7 | HISAT2 | GRCh38 | Ensembl | 40.0% | 16394 | 41.7% | 16653 | 0 | 0 | 7 | 0 | 7 |
| HS7 | HISAT2 | T2T | Ensembl | 38.3% | 15767 | 39.8% | 15975 | 0 | 0 | 7 | 0 | 7 |
| HS7 | STAR | GRCh38 | Ensembl | 41.2% | 16683 | 49.0% | 16835 | 0 | 1 | 6 | 0 | 7 |
| HS7 | STAR | T2T | Ensembl | 39.8% | 16092 | 47.3% | 16162 | 0 | 5 | 2 | 0 | 7 |
| Summary |  |  | Ensembl | 36.6% | 14716 | 41.4% | 15599 | 0 | 10 | 58 | 0 | 68 |

Users may choose the most suitable assembler or pipeline according to their specific requirements. For examples, if a more comprehensive assembly is preferred, particularly when transcripts with retained introns are needed, then Scallop2 may be the best choice among Scallop2-ft, StringTie2, and StringTie2-ft; on the other hand, if transcripts with retained introns are not needed, then Scallop2-ft would be the best option among the other choices.

## 3 Conclusions

In this work we assess the impact of annotations on transcript assembly. We have discovered that different conclusions can arise when using different annotations for evaluation. To unravel this mystery, we analyzed the transcript biotypes across various annotations and assemblies, and figured that the bias in annotating and assembling transcripts with retained introns is the main cause. Moreover, we investigated similarities and differences in annotations at the intron-exon boundary, junction, and intron-chain levels, and our results indicate that the primary structural divergences in annotations occur at the intron-chain level.

In addition, we have developed a standalone tool for extracting or filtering out transcripts with retained introns from an assembly, freely available at https://github.com/Shao-Group/irtool. This tool can be used in conjunction with any assembler to mitigate bias in transcript assembly with intron retentions. Our results show that the accuracy improvement varies significantly when applying this tool to different assemblers. Specifically, we found that Scallop2-ft (Scallop2 followed by filtering) is superior to Scallop2, StringTie2, and StringTie2-ft when producing an assembly without intron-retained transcripts.

## 4 Methods

### 4.1 Evaluation Pipeline

We compare two widely-used reference-based assemblers, StringTie2 (version 2.2.0) and Scallop2 (version 1.1.2), both of which are tested with their default parameters in all experiments. We use two human RNA-seq datasets: EN10, consisting of 10 paired-end RNA-seq samples downloaded from the ENCODE project, and HS7, containing 7 paired-end RNA-seq samples used in the Long Read Genome Annotation Assessment Project. The assembled transcripts are assessed using five transcriptome annotations derived from two human genome builds: GRCh38 and T2T-CHM13. GRCh38 is the most commonly used human genome assembly, while T2T-CHM13 is the most recent and comprehensive, gapless sequence of a human genome. For GRCh38, we use the latest Ensembl annotation (release 107 on genome build GRCh38.p13) and RefSeq annotation (release 110 on genome build GRCh38.p14). We use 3 annotations from T2T-CHM13, namely the Ensembl annotation, RefSeq annotation, and its own CHM13 annotation released with its paper.

We use the pipeline depicted in Figure 11 to assess the two assemblers’ accuracy using different annotations. Each RNA-seq sample is aligned with two popular splice-aware aligners, STAR [33] and HISAT2 [34]. The resulting read alignments will be piped to the assemblers, producing a set of assembled transcripts.

**Figure 11:**
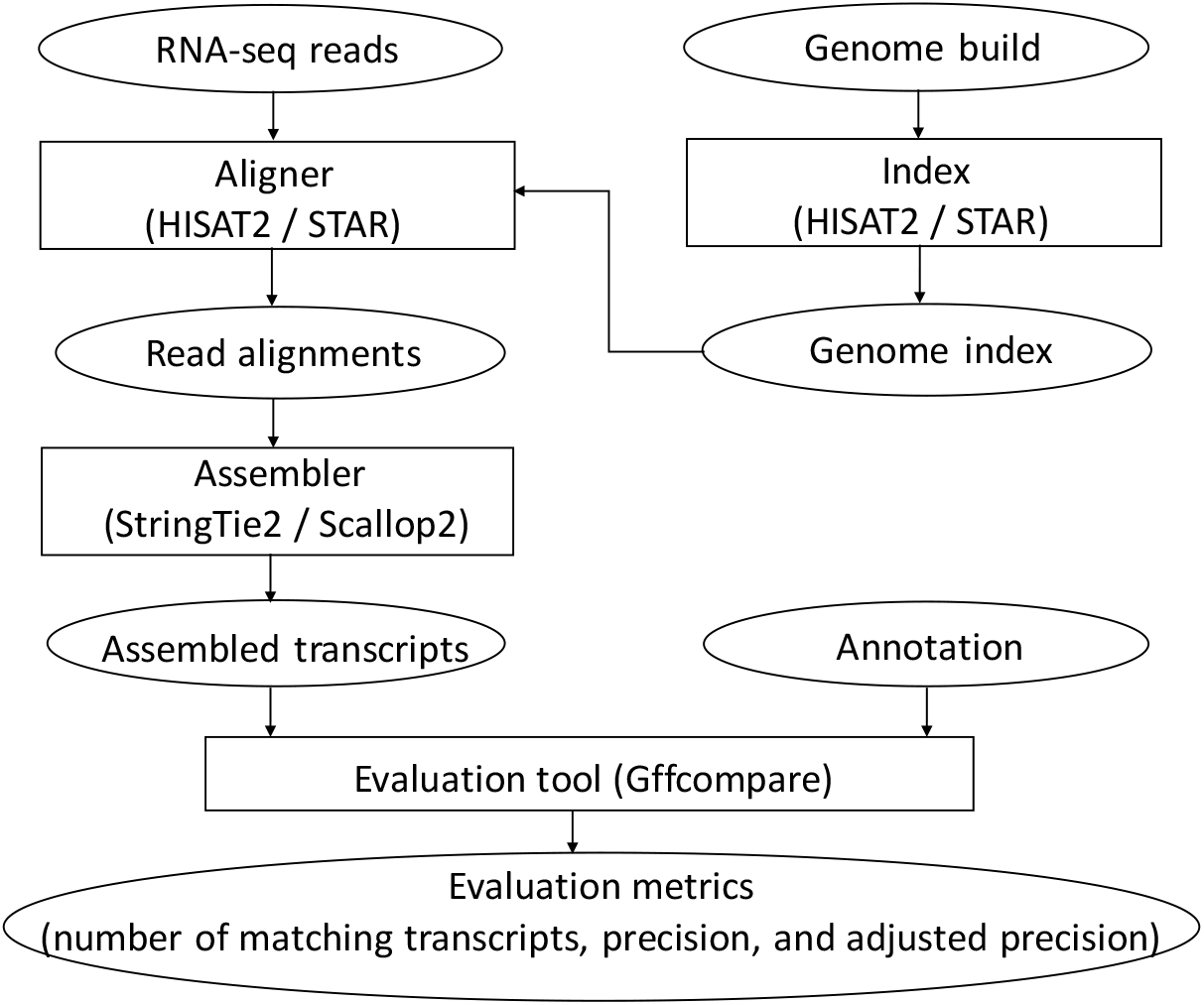
Pipeline of evaluating the accuracy of compared assemblers.

The accuracy of the assembled transcripts will be evaluated using tool gffcompare [35], with one of the annotations serving as the ground-truth. We use the “transcript level” measure defined by gffcompare: an assembled multi-exon transcript is considered to be “matching” if its intron-chain exactly matches that of a transcript in the annotation; an assembled single-exon transcript is defined as “matching” if there is a significant overlap (80% by default) with a single-exon transcript in the annotation. We report the two metrics calculated by gffcompare: the total number of matching transcripts, which is proportional to recall, and precision, defined as the total number of matching transcripts divided by the total number of assembled transcripts. On samples where mixed outcomes exhibit, i.e., one method performs better on one metric but not on the other, we compare their *adjusted precisions*, defined as their precisions when matching transcripts are adjusted to be the same. Specifically, we take the assembly with higher recall and gradually remove its transcripts with lowest (predicted) abundance. In this process its recall will drop but its precision will likely increase as abundance is highly correlated with accuracy. We step this process when its recall matches that of the other assembler, and the precision at this moment (i.e., the adjusted precision) will be compared with the precision of the other assembler. This way of comparison has been used in previous studies [28, 30].

### 4.2 Metrics for Structural Similarities

We propose a set of metrics for measuring the structural similarities of two annotations. Let *t* be a transcript. We use *B*(*t*), *J*(*t*), and *C*(*t*) to represent the set of intron-exon boundaries, the set of junctions, and the intron-chain, of *t*, respectively. Let *g* be a gene, which may contain multiple transcripts in an annotation, we define *B*(*g*) = ∪ *_t_*_∈*g*_*B*(*t*), *J*(*g*) = ∪*_t_*_∈*g*_*J*(*t*), and *C*(*g*) = ∪*_t_*_∈*g*_*C*(*t*), to represent the set of boundaries, junctions, and intron-chains, of all transcripts in gene *g*, respectively. Let *T* be an annotation with many annotated genes. We then define *B*(*T*) = ∪*_g_*_∈*T*_ *B*(*g*), *J*(*T*) = ∪*_g_*_∈*T*_ *J*(*g*), and *C*(*T*) = ∪*_g_*_∈*T*_*C*(*g*). We use *Jaccard similarity* to measure the structural similarity of two annotations. Formally, let *T*_1_ and *T*_2_ be two annotations, we use *J_B_*(*T*_1_*, T*_2_) := |*B*(*T*_1_) ∩ *B*(*T*_2_)|*/*|*B*(*T*_1_) ∪ *B*(*T*_2_)|, *J_J_*(*T*_1_*, T*_2_) := |*J*(*T*_1_) ∩ *J*(*T*_2_)|*/*|*J*(*T*_1_) ∪ *J*(*T*_2_)|, and *J_C_*(*T*_1_*, T*_2_) := |*C*(*T*_1_) ∩ *C*(*T*_2_)|*/*|*C*(*T*_1_) ∪ *C*(*T*_2_)|, to measure the similarity of *T*_1_ and *T*_2_ at the level of boundary, junction, and intron-chain, respectively. Please see Figure 12 for an illustration of these definitions.

**Figure 12:**
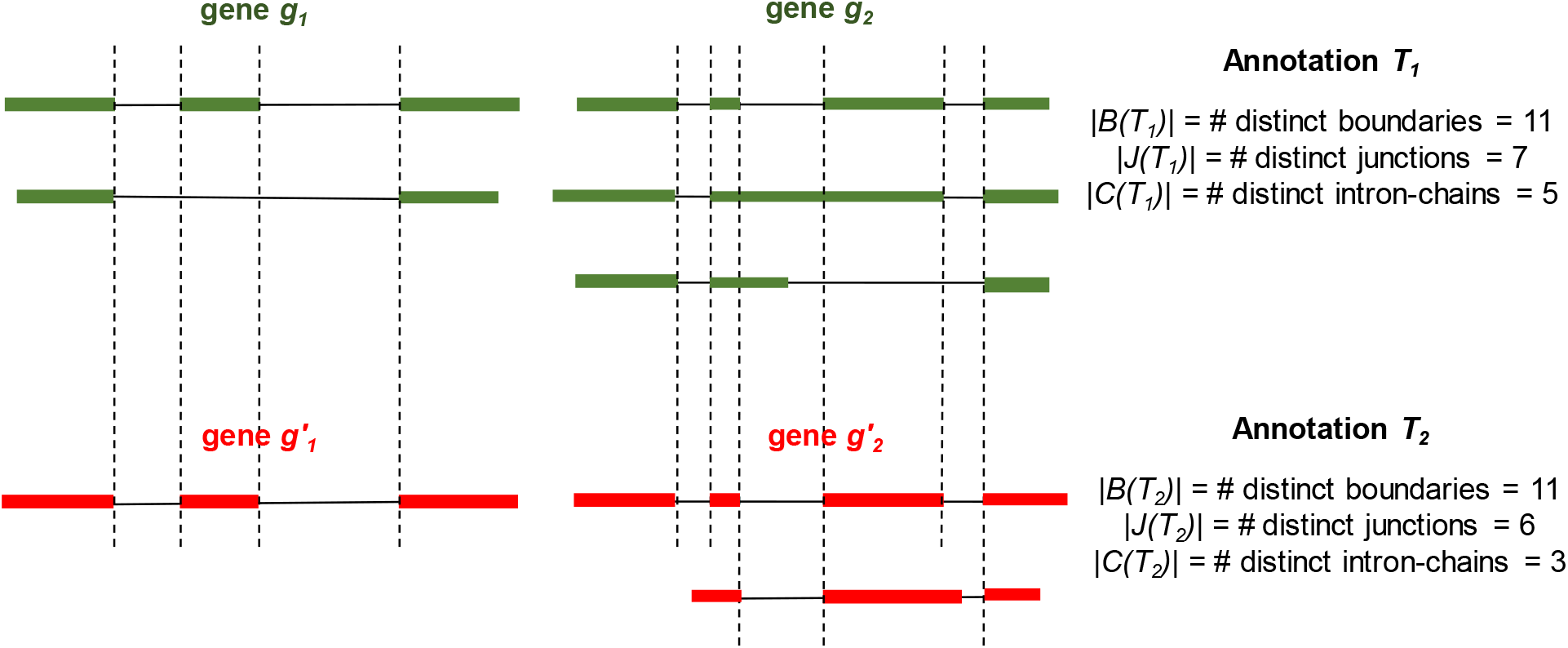
A toy example for illustrating the Jaccard similarity of two annotations *T*_1_ and *T*_2_ at the level of boundary, junction, and intron-chain. Genes and transcripts from the same annotation are colored the same. Identical boundaries between two annotations are marked with vertical dashed lines. We have *J_B_*(*T*_1_*, T*_2_) := |*B*(*T*_1_) ∩ *B*(*T*_2_)|*/*|*B*(*T*_1_) ∪ *B*(*T*_2_) = 5*/*6|, *J_J_*(*T*_1_*, T*_2_) := |*J*(*T*_1_) ∩ *J*(*T*_2_)|*/*|*J*(*T*_1_) ∪ *J*(*T*_2_)| = 5*/*8|, and *J_C_*(*T*_1_*, T*_2_) := |*C*(*T*_1_) ∩ *C*(*T*_2_)|*/*|*C*(*T*_1_) ∪ *C*(*T*_2_)| = 1*/*3|.

### 4.3 Constructing Gene Correspondence

We focus on multi-exon genes when constructing the correspondence between genes in two annotations, i.e., genes annotated with at least one multi-exon transcript. We propose a simple approach: two genes *g*_1_ ∈ *T*_1_ and *g*_2_ ∈ *T*_2_, where *T*_1_ and *T*_2_ are two annotations, form a pair if they share at least one intron-exon boundary, i.e., 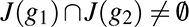. Note that in this definition one gene in *T*_1_ may form gene pairs with multiple genes in *T*_2_, but this rarely happens since two genes in one annotation normally do not share intron-exon boundaries. The Jaccard similarity of each constructed gene pair (*g*_1_*, g*_2_) at boundary, junction, and intron-chain levels, can be defined similarly, formally written as *J_B_*(*g*_1_*, g*_2_) := |*B*(*g*_1_) ∩ *B*(*g*_2_)|*/*|*B*(*g*_1_) ∪ *B*(*g*_2_)|, *J_J_*(*g*_1_*, g*_2_) :=|*J*(*g*_1_) ∩ *J*(*g*_2_)|*/*|*J*(*g*_1_) ∪ *J*(*g*_2_)|, and *J_C_*(*g*_1_*, g*_2_) := |*C*(*g*_1_) ∩ *C*(*g*_2_)|*/*|*C*(*g*_1_) ∪ *C*(*g*_2_)|.

### 4.4 Definition of Intron Retentions in the Context of Assembly

We describe our definition of (partial) intron retentions. To determine if a transcript *t* in an assembly has intron retention or not, we need to find another transcript *r* in the same assembly as reference, and compare *t* with *r*. The definition also uses the abundances (i.e., expression levels) of *t* and *r*; we therefore assume that each transcript in the assembly are associated with an abundance. Most assemblers, including StringTie2 and Scallop2, assembles transcripts while also predicting their abundances. Let *p*(*t*) and *p*(*r*) be the abundances of transcripts *t* and *r*, respectively. We define transcript *t* has *intron retention* if there exists transcript *r* (in the same assembly with *t*) such that *p*(*r*)*/p*(*t*) is above a threshold (0.5 by default) and either (a) the first exon of *t* spans an intron and the following exon of *r* (Figure 13, criterion 1), or (b) the last exon of *t* spans an exon and the following intron of *r* (Figure 13, criterion 2), or (c) there exists an exon in *t* and an intron in *r* such that the intron is fully covered by the exon (Figure 13, criterion 3). Note that one transcript *t* may satisfy two or more criteria with the same *r*, or may satisfy one or more criteria with different *r*.

**Figure 13:**
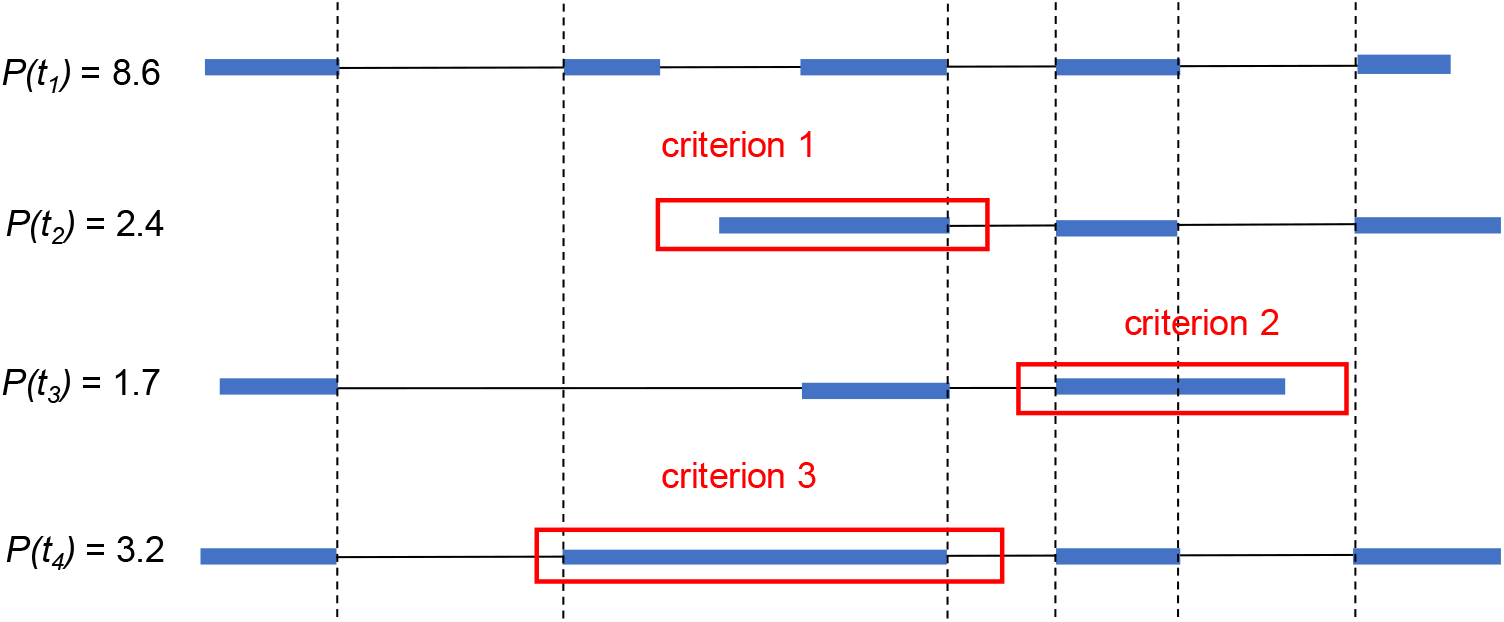
An illustrative example for the three criteria used to define transcripts with intron retentions. Identical boundaries are marked with vertical dashed lines. Transcript *t*_2_ satisfies criterion 1 (it has lower abundance than *t*_1_, and its first exon spans the second intron of *t*_1_). Transcript *t*_3_ satisfies criterion 2 (it has lower abundance than *t*_1_ and its last exon spans the fourth intron of *t*_1_). Transcript *t*_4_ satisfies criterion 3 (it has lower abundance than *t*_1_ and its second exon fully covers the second intron of *t*_1_).

## Declarations

### Availability of data and materials

The EN10 dataset used in this manuscript are downloaded from the ENCODE project [13], consisting of 10 paired-end RNA-seq samples. Their SRA accession IDs are SRR307903, SRR307911, SRR315323, SRR315334, SRR387661, SRR534291, SRR534307, SRR534319, SRR545695, and SRR545723. The HS7 dataset consists of 7 paired-end RNA-seq samples downloaded from the Long Read Genome Annotation Assessment Project. Their accession IDs are ENCFF766OAK paired with ENCFF644AQW, ENCFF198RQU paired with ENCFF620HBM, ENCFF247XJT paired with ENCFF785SWH, ENCFF201EVI paired with ENCFF591ISP, ENCFF221SLD paired with ENCFF223VFL, ENCFF145IIO paired with ENCFF597GZT, and ENCFF701OIK paired with ENCFF139HIY.

### Competing interests

The authors declares no competing interests.

### Funding

This work is supported by the US National Science Foundation (DBI-2019797 and DBI-2145171 to M.S.) and by the US National Institutes of Health (R01HG011065 to M.S.).

## Notes

### Competing Interest Statement

The authors have declared no competing interest.

